# Neural correlates of novel word-form learning in developmental language disorder

**DOI:** 10.64898/2026.03.28.715039

**Authors:** Nilgoun Bahar, Gabriel J. Cler, Salomi S. Asaridou, Harriet J. Smith, Hanna E. Willis, Máiréad P. Healy, Sana Chughtai, Meron Haile, Saloni Krishnan, Kate E. Watkins

**Author notes:** **Corresponding author:** Nilgoun Bahar, UCSF Dyslexia Center, Sandler Neurosciences Center, 675 Nelson Rising Lane, Suite 190, San Francisco, California., **Email:**. Joint senior authors (Saloni Krishnan, Kate E. Watkins).

## Abstract

Children with developmental language disorder (DLD) have persistent language learning difficulties and often perform poorly on pseudoword repetition, a task that probes phonological, memory, and speech-motor processes that support vocabulary acquisition. Research on the neural basis of pseudoword repetition in DLD is limited. We used whole-brain functional MRI (fMRI) to examine pseudoword repetition and repetition-based learning in 46 children with DLD (ages 10–15 years) and 71 age-matched children with typical language development. During scanning, children heard and repeated pseudowords paired with visual referents, allowing us to track learning-related changes in neural activity across repetitions. Repeated pseudoword production yielded comparable behavioural learning across groups, with faster productions by later repetitions. Post-scan, form–referent recognition was comparable across groups, whereas pseudoword repetition accuracy was lower in DLD. Pseudoword repetition engaged a distributed neural network, including inferior frontal cortex bilaterally (greater on the left), premotor and sensorimotor cortex, and posterior temporal and occipital regions. Group differences emerged primarily in regions where activity was task negative (i.e., below baseline or deactivated): lateral occipito-parietal cortex (posterior angular gyrus), medial parieto-occipital cortex (retrosplenial), and right posterior cingulate cortex. Learning-related decreases in activity were similar across groups, but region-of-interest analyses showed reduced leftward lateralisation of activity in inferior frontal gyrus in DLD. These findings suggest weaker disengagement of the default mode network during a linguistically demanding task in DLD. Although repetition-based pseudoword learning recruited similar neural mechanisms in both groups, these mechanisms may operate less efficiently in DLD, alongside reduced hemispheric specialisation in inferior frontal cortex.

**Highlights:**

1. Similar repetition-related neural attenuation across groups during pseudoword learning.
2. Reduced default-mode network suppression during pseudoword repetition in DLD.
3. Reduced left-hemisphere specialisation of inferior frontal cortex in DLD.
4. Repetition-based learning in DLD supported by less efficient neural networks.

## 1. INTRODUCTION

Learning new words is a key component of language acquisition, allowing us to communicate thoughts, emotions, and ideas with increasing clarity. Most children learn new words quickly, forming robust links between novel word forms and their meanings after only a few exposures (Carey & Bartlett, 1978). Developmental language disorder (DLD) is a neurodevelopmental condition characterised by persistent difficulties in understanding and producing language that cannot be attributed to hearing loss, intellectual disability, or other biomedical causes (Bishop et al., 2016, 2017). For children with DLD, word learning is often gradual, requiring more repetition or support than their peers with typical language development (Jackson et al., 2019; Rice & Hoffman, 2015). These word learning differences appear to arise primarily from difficulties with encoding new word forms in this population, whereas mapping word forms onto referents (i.e., word recognition) is a relative strength (Bishop & Hsu, 2015; Kan & Windsor, 2010; Leonard et al., 2021; McGregor et al., 2017). This profile is perhaps best reflected in pseudoword repetition. Children with DLD often perform poorly on pseudoword repetition tasks compared with their peers with typical language, and these differences tend to persist into adulthood even when broader language difficulties become less apparent (Bishop et al., 1996; Schwob et al., 2021). In this paper, we examined brain activity in children with and without DLD during pseudoword repetition to gain insight into the neural processes that support word learning.

Accurately repeating a pseudoword requires holding a novel phonological sequence in memory and coordinating its corresponding motor pattern (Coady & Evans, 2008; Gathercole et al., 2006; Krishnan et al., 2013). The ability to repeat pseudowords is a strong predictor of vocabulary size (Adlof & Patten, 2017; Gathercole et al., 1999; Hoff et al., 2008). Difficulties with pseudoword repetition, together with evidence of reduced automatisation of non-linguistic sequences in DLD, have led some researchers to propose that language difficulties in this population reflect a broader impairment in learning and integrating complex sequences regardless of modality (Goffman & Gerken, 2023; Krishnan et al., 2016; Ullman et al., 2020). According to the Procedural Deficit Hypothesis (Ullman & Pierpont, 2005) and its recent extension, the Procedural circuit Deficit Hypothesis (Ullman et al., 2020), individuals with DLD have disruptions in cortico-basal ganglia-thalamo-cortical circuits, networks that support implicit learning processes fundamental to learning sequential patterns, including novel phonological forms. Supporting this, we previously found structural brain differences in the inferior frontal cortex (Bahar et al., 2024) and dorsal striatum (Cler et al., 2026; Krishnan et al., 2021) in children with DLD. A recent meta-analysis by Ullman and colleagues (2024) reported that neural differences in DLD are most consistently observed in the basal ganglia, with structural differences in the anterior neostriatum identified in 100% of the participant groups in which this region was examined, and functional differences reported in 79% of them.

Studying the neural basis of pseudoword repetition could provide a window into the neurocognitive mechanisms postulated by the Procedural (circuit) Deficit Hypothesis. In adults with typical language abilities, pseudoword repetition engages a distributed network encompassing posterior superior temporal and supramarginal regions, portions of the medial and lateral frontal cortex, and subcortical structures including the basal ganglia, thalamus, and cerebellum (Rauschecker et al., 2008; Simmonds et al., 2014; Tettamanti et al., 2005). Yet, evidence for atypical engagement of these circuits in DLD is limited and mixed. One fMRI study of overt pseudoword repetition in 9- to 11-year-olds (*n* = 18) found no reliable neural differences between DLD and typically developing peers, despite weaker performance in the DLD group on the pseudoword repetition task (Pigdon et al., 2020). In contrast, structural differences and pronounced underactivation in cortical motor areas and subcortical regions (including the putamen and cerebellum) were observed in affected members of the KE family, whose speech and language difficulties could be considered as more severe cases of DLD (Argyropoulos et al., 2019; Liégeois et al., 2011). These findings suggest that neural differences may emerge more clearly in individuals with greater impairment severity. Supporting this, Pigdon and colleagues (2020) noted that their participants with DLD had relatively mild deficits compared with other samples.

Understanding neural changes during pseudoword *learning* may also provide further insight into mechanisms that differ in DLD. Repeated exposure to pseudowords enhances their phonological and articulatory representations; with practice and increasing familiarity, performance tends to become more efficient and automatised, potentially drawing on striatal circuitries that support sequence learning, habit formation, and detecting statistical regularities of input (Graybiel, 2008; Orpella et al., 2021; Packard & Knowlton, 2002; Turk-Browne et al., 2009). Rauschecker and colleagues (2008) used a repetition paradigm in which typically developing adults were asked to covertly repeat pseudowords, some of which were presented four times without participants’ awareness. Participants pressed a button to indicate when each covert repetition was complete. The authors found that covert repetition durations decreased with repeated exposure. Neurally, repetition was associated with decreased activity across left-lateralised temporal and frontal regions linked to language processing, suggesting more efficient engagement of these regions as speech motor patterns became more practised (Rauschecker et al., 2008). To date, only one fMRI study has examined a roughly similar phenomenon of language learning in DLD. Plante and colleagues (2017) assessed implicit learning of syllable sequences representing novel word forms in young adults with DLD (*n* = 16) and found more widespread activity than in controls in inferior frontal and superior temporal cortex, as well as the supramarginal gyrus.

In this study, we used fMRI to compare brain activity during pseudoword repetition and changes in brain activity during pseudoword learning in 46 10- to 15-year-olds with DLD and 71 age-matched controls. We distinguished between (i) participants’ initial productions of pseudowords (first-time repetitions), which served as a proxy for phonological short-term memory and speech-motor skills, and (ii) changes over three subsequent repetitions, which served as a measure of implicit learning (without explicit feedback on performance). Based on previous findings in typically developing adult learners (Rauschecker et al., 2008), we anticipated learning-related decreases in brain activity across cortical regions involved in speech and language. We also expected involvement of subcortical regions implicated in learning, consistent with accounts proposing a role for cortico-striatal systems in language learning (Orpella et al., 2021) and its implications in DLD (Ullman & Pierpont, 2005, Ullman et al., 2020). We expected to see behavioural and neural differences between typically developing and DLD groups during this learning process.

## 2. METHODS

### 2.1 Ethics

Ethical approval was granted by the University of Oxford’s Medical Sciences Interdivisional Research Ethics Committee (R55835/RE002). Written parental informed consent and child assent were obtained prior to participation.

### 2.2 Participants

#### Recruitment

A total of 175 participants aged 10–15 years took part in the Oxford Brain Organisation in Language Development (OxBOLD) project (see Asaridou et al., 2024; Bahar et al., 2024; Cler et al., 2026; Krishnan et al., 2021, 2022). All participants had been raised in the UK learning English before age five. Recruitment took place through social media, community and school outreach, specialist organisations, prior research databases, and local advertising. Typically developing controls were primarily recruited through community advertisements.

Parents first completed a telephone screening that included the Strengths and Difficulties Questionnaire (SDQ; Goodman, 1997) and the Social Communication Questionnaire (SCQ; Rutter et al., 2003). The SDQ was used to identify elevated behavioural difficulties (particularly hyperactivity and inattention symptoms relevant to ADHD). The SCQ screened for autistic traits. Parents also completed an intake questionnaire and the Children’s Communication Checklist-2 (CCC-2; Bishop, 2003) to provide demographic and social communication information. Children were excluded if parents reported (1) SDQ Hyperactivity score above seven or SCQ score above 15; (2) a known neurological, genetic, or other developmental condition; (3) vision or hearing impairment; or (4) MRI contraindications. Left handedness and multilingualism were not exclusion criteria.

#### DLD and TD criteria

Participants who passed the screening were invited for in-person testing to complete a comprehensive neuropsychological test session assessing language and other cognitive and motor skills. All passed a bilateral pure-tone audiometric screening. Following assessment, participants were classified as having DLD if they had a documented history of speech-language difficulties and scored ≥ 1 standard deviation (*SD*) below the mean on at least two of six standardised language measures assessing receptive and expressive vocabulary, grammar, and narrative skills (see Supplementary Table 1 for a list of tests and group results), in line with established diagnostic criteria (Hsu & Bishop, 2014; Tomblin et al., 1996). Typically developing participants (TD) had no such history and no more than one language score ≥ 1 *SD* below the mean.

#### Final sample

Of the 175 participants recruited, data from 58 were excluded for the following reasons: not raised in the UK with English as the primary language before age five (identified after testing rather than at screening; *n* = 3); incomplete behavioural assessment (*n* = 2); a history of speech-language difficulties but not meeting DLD criteria at assessment (*n* = 28); poor language performance despite no reported history of speech-language difficulties (*n* = 1); nonverbal intelligence quotient (IQ) < 70, defined as a score ≥ 2 *SD*s below the mean on a composite of the Block Design and Matrix Reasoning subtests of Wechsler Intelligence Scale for Children– Fourth Edition WISC-IV; Wechsler, 2004) (*n* = 3); incomplete MRI (*n* = 3); incidental MRI findings (*n* = 3); or excessive head motion during the fMRI task (> 2.4 mm; *n* = 15).

The final sample comprised 117 participants with complete neuropsychological and fMRI data: 46 with DLD and 71 TD controls. The groups were well matched in age (TD: *M* = 12.6 years, *SEM* = 0.2; DLD: *M* = 12.4 years, *SEM* = 0.3) and showed comparable age ranges (TD: 10;0– 15;7; DLD: 10;0–15;11). They also did not differ significantly in sex distribution (TD: 56% male, *n* = 40; DLD: 70% male, *n* = 32), χ^2^(1) = 1.54, p = .214, or handedness (TD: 86% right-handed, *n* = 61; DLD: 83% right-handed, *n* = 38), χ^2^(1) = 0.04, p = .824.

### 2.3 In-scanner task

During the scan, participants were instructed to overtly repeat pseudowords played through noise-cancelling headphones while viewing corresponding images of ‘aliens’ (Fig 1A for a schematic of the experiment). Participants were not informed that some stimuli would repeat.

**Fig 1.**
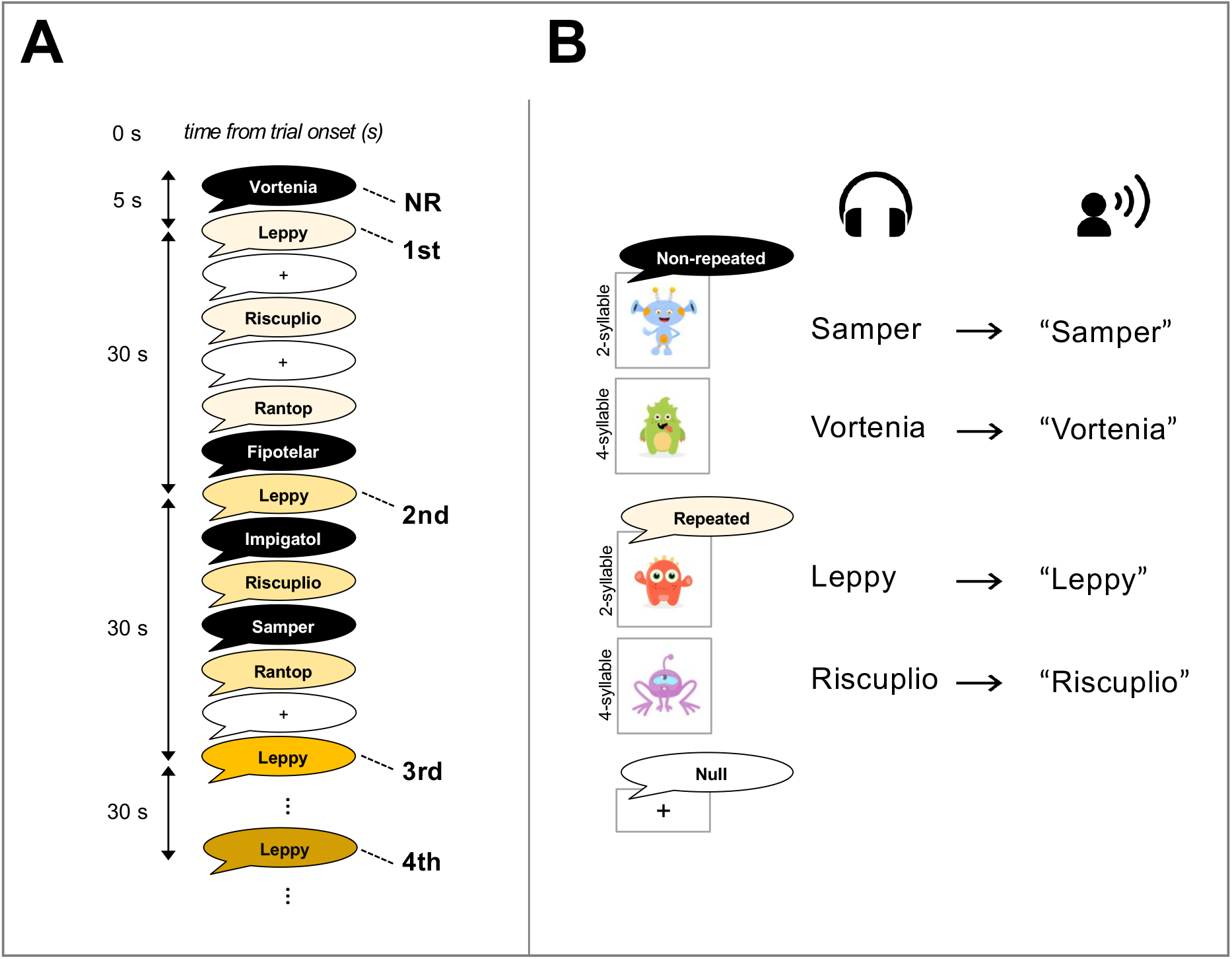
Schematic of experimental design. (**A**) Half of the items were presented only once (NR: non-repeated; black background), while the remaining items were presented four times (1st–4th presentations; yellow gradient from light to dark). Colours are shown for illustration only, participants did not see colour in the scanner. Blank screens with a fixation cross represent null trials with no stimulus. (**B**) On each trial, participants heard a pseudoword through noise-cancelling headphones and were instructed to repeat it aloud while a corresponding alien image appeared on the screen. Both the repeated and non-repeated conditions contained 16 stimuli, each with an equal number of 2- and 4-syllable items.

#### Stimuli

We generated 32 phonotactically plausible English pseudowords using the Wuggy Pseudoword Generator (Keuleers & Brysbaert, 2010; Appendix A). Half of the pseudowords (*n* = 16) had two syllables (e.g., “Samper”), and the other half (*n* = 16) had four syllables (e.g., “Vortenia”). A native British-English female speaker recorded all items. For form-referent learning, each pseudoword was deterministically paired with a unique, visually distinct ‘alien’ image selected from the <u>Shutterstock repository</u> (Fig 1B).

#### In-scanner task design

Half of the pseudowords were repeated four times at 30-sec intervals, intermixed with 16 pseudowords presented only once (Fig 1A). This resulted in 80 total stimulus presentations (16 repeated × four + 16 non-repeated). Stimuli were presented every five seconds, with randomly inserted null events (fixation cross) also five-sec duration. Task compliance was monitored in real time by researchers observing participants’ overt responses.

### 2.4 MRI acquisition

Participants completed functional and structural scans as part of an hour-long scan session (see Krishnan et al., 2021). The pseudoword repetition task followed a verb generation task and was conducted on a 3T Siemens Prisma scanner with a 32-channel head coil. Participants wore earplugs and Optoacoustics noise-cancelling headphones (up to 30 dB attenuation), with foam and inflatable pads to minimise movement. Stimuli were clearly audible and overt responses were recorded using an Optoacoustics FOMRI-III microphone.

Functional imaging closely followed ABCD study task fMRI parameters: 625 T_2_*-weighted EPI volumes (TR = 800 ms, TE = 30 ms, flip angle = 52°, FOV = 216 mm, multiband factor = 6), with 2.4 mm^3^ resolution. The first 25 volumes were discarded to allow the noise-cancelling system to stabilise; the remaining 600 volumes were used. B0 field maps were also collected. Structural images were acquired using a T_1_-weighted magnetisation-prepared rapid acquisition gradient echo sequence (MP-RAGE; TR = 1900 ms, TE = 3.97 ms, flip angle = 8°, TI = 904 ms) with 1-mm isotropic resolution over five minutes, 30 seconds.

### 2.5 MRI analysis

#### Pre-processing

Functional imaging data were analysed using the General Linear Model in FEAT v6.0.0 (FSL; Smith et al., 2004). Preprocessing included motion correction (MCFLIRT; Jenkinson, 2002), skull stripping (BET; Smith, 2002), and 6-mm FWHM spatial smoothing. Data were mean-normalised and temporally high-pass filtered (90-sec cut-off) to remove low-frequency drift. Field maps were generated from unwrapped phase maps and used to correct susceptibility distortions. Functional scans were aligned to each participant’s T_1_-weighted structural image using boundary-based registration (Greve & Fischl, 2009), and then transformed to MNI-152 standard space using FNIRT (FNIRT; Andersson et al., 2008).

#### First-level analysis

The first-level model included four explanatory variables (EVs): the first presentation of both repeated and non-repeated pseudowords (1st), followed by the second (2nd), third (3rd), and fourth (4th) presentations of repeated pseudowords. Including both repeated and non-repeated trials in the first EV enabled more reliable effect size estimation. Each EV was modelled with a duration of five seconds to capture both auditory perception and overt speech. Null events were not modelled explicitly but contributed to the implicit baseline. Temporal derivatives were included for all EVs, and six motion parameters (translations and rotations in X, Y, and Z) were entered as nuisance regressors. Haemodynamic responses were modelled using a double-gamma haemodynamic response function convolved with stimulus timing. Primary contrasts compared the 1st, 2nd, 3rd, and 4th presentations against implicit baseline. Learning-related changes across repeated items were assessed using linear decrease contrasts across the repeated presentations.

#### Group-level analysis

Group averages and between-group comparisons (TD vs. DLD) were conducted using FMRIB’s Local Analysis of Mixed Effects (FLAME 1; Beckmann et al., 2003; Woolrich et al., 2004). Results were cluster-corrected using Gaussian Random Field Theory with a Z > 3.1 threshold and family-wise error correction at *p* < .05.

We additionally created probabilistic overlap maps to evaluate the consistency of BOLD activity across each group. For each participant, we determined voxel-wise significance thresholds based on data smoothness and calculated the number of resolution elements (RESELs) by dividing the whole-brain volume by RESEL size. Using Gaussian Random Field maximum height theory and the number of RESELs, we calculated the corrected voxel-wise Z score significant at *p* = .05. Each participant’s map was thresholded at this Z value, binarised and registered to MNI space. The binarised maps were summed and divided by group size, resulting in overlap maps showing the proportion of participants with significant activity at each voxel.

#### ROI analysis

In a separate post hoc analysis, percent signal change from baseline during the first and subsequent pseudoword repetitions was extracted from nine regions of interest (ROIs) using the Harvard-Oxford Cortical and Jülich Histological atlases. These ROIs were selected as part of the cortico-striatal circuitry, connecting cortical-subcortical networks involved in implicit learning and skill acquisition. ROIs included pars triangularis, pars opercularis, caudate nucleus, and putamen all bilaterally, and one that included both left and right supplementary motor area (since these are in the midline). Masks from the atlases were thresholded at 30%, binarised, and applied to each participant’s functional data to extract median signal change in these regions for each presentation of the repeated pseudowords.

Linear mixed-effects models tested neural activity patterns, with fixed effects for repetition trial (1st–4th), group (TD vs. DLD), hemisphere (left vs. right; except for the analysis of the supplementary motor area), and their interactions, with random intercepts for repetition trials and participants. The 1st trial, TD group, and left hemisphere were set as reference levels. A Bonferroni-corrected significance threshold of *p* = .01 was applied for five tests (five ROIs). These analyses were performed using the lme4 package (Bates et al., 2015) in R v4.3.2 (R Core Team, 2023).

### 2.6 Behavioural data

Two sets of behavioural data were collected: in-scanner and post-scan. Because in-scanner audio recordings were noisy and time-intensive to code, analyses were conducted on a randomly selected subset of participants, approximately one third of the sample (17 TD, 17 DLD). Two research staff, blind to group membership, independently coded pseudoword repetitions from each repetition trial (trials 1–4), scoring repetition accuracy (correct/incorrect) and measuring pseudoword production duration (i.e., stimulus onset latency to the end of the overt response). Disagreements were resolved by discussion until consensus was reached.

Post-scan behavioural testing took place approximately one hour after the in-scanner task. Participants first repeated all 32 pseudowords aloud, with responses audio-recorded for offline scoring, and then completed a three-alternative forced-choice (3AFC) pseudoword–picture recognition task. Due to technical issues, post-scan repetition data were unavailable for three TD and two DLD participants, yielding a final sample of 68 TD and 44 DLD for repetition accuracy analyses. Recognition data were missing for one TD and one DLD participant, resulting in a final sample of 70 TD and 45 DLD. Two research staff independently scored post-scan repetition accuracy; disagreements (8.68% of 4,158 trials) were resolved through discussion, with remaining cases adjudicated by a third experimenter.

Behavioural data were analysed using mixed-effects models. In-scanner production duration was analysed with linear mixed-effects models, and in-scanner repetition accuracy with generalised linear mixed-effects models with a logit link. Both models included fixed effects of trial (1–4), group (TD vs. DLD), and pseudoword length (2 vs. 4 syllables), with random intercepts for participants; reference levels were trial 1, TD group, and 2-syllable pseudowords. Post-scan recognition and repetition accuracy were analysed using generalised linear mixed-effects models with fixed effects of presentation type (repeated vs. non-repeated), group, and pseudoword length, again with random intercepts for participants. Reference levels were non-repeated pseudowords, TD group, and 2-syllable pseudowords. In all analyses, we first assessed interactions, and in the absence of a significant interaction, we looked at the main effects.

## 3. RESULTS

### 3.1 Behavioural data

#### In-scanner pseudoword learning

Repeated presentation of pseudowords for overt repetition in the scanner resulted in a behavioural learning effect in both groups (Fig 2). There were no significant interactions between repetition trials, group, and pseudoword length (all *p*s > .23). Across groups and pseudoword lengths, the duration of pseudoword productions decreased with repetition, showing significantly shorter duration by the 4th trial compared with the 1st trial (β = −0.094, SE = 0.029, CI [−0.15, −0.03], *p* = .001; Fig 2A). The durations were not statistically different between groups averaged across repetition trials and syllable lengths (main effect of group, *p* = .969). The durations were significantly longer for 4-syllable than for 2-syllable pseudowords across groups and repetition trials (main effect of length, β = 0.441, SE = 0.024, CI [0.39, 0.48], *p* < .001).

**Fig 2.**
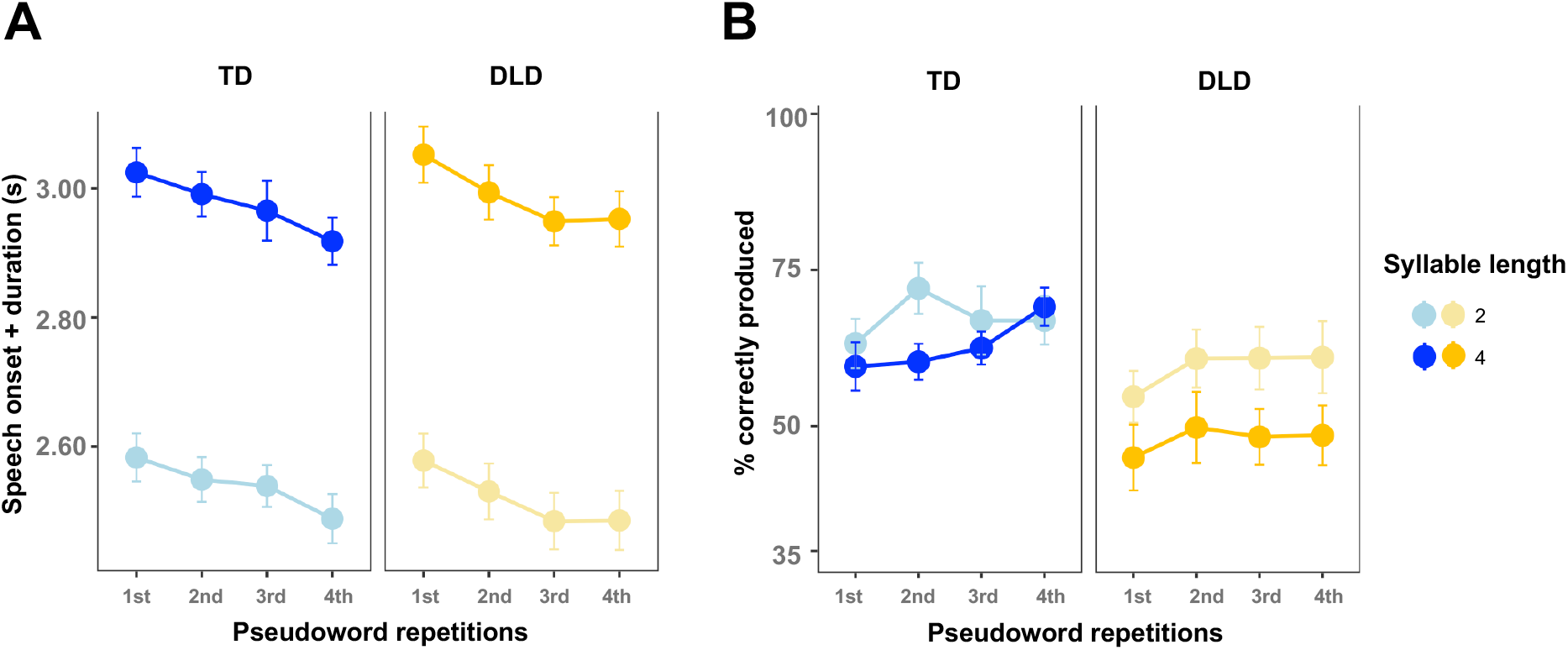
Behavioural performance during in-scanner overt pseudoword repetition. (**A**) Pseudoword production duration (stimulus onset latency + duration of overt response) in seconds and (**B**) percentage of accurately produced pseudowords per trial are shown across four stimulus repetitions for typically developing (TD) and developmental language disorder (DLD) groups, separately for 2- and 4-syllable pseudowords. Error bars represent standard errors of the mean (*SEM*).

The interactions between repetition trials, group, and pseudoword length for pseudoword production accuracy were not significant (all *p*s > .22; Fig 2B). Across repetitions, pseudoword production accuracy was consistently above 50% in the TD group and for 2-syllable stimuli in the DLD group but remained near 50% for 4-syllable stimuli in the DLD group. However, these apparent differences were not statistically reliable: there was no significant main effect of group (*p* = .159), or pseudoword length (*p* = .370). There was a trend toward increased accuracy from the 1st to the 2nd repetition across groups and pseudoword lengths (*p* = .071), with no further improvement on the 3rd or 4th repetitions relative to the 1st repetition trial (both *p*s = .457).

#### Post-scan pseudoword recognition and repetition accuracy

We assessed how accurately participants recognised the images associated with each pseudoword, focusing on the effects of presentation type, group, and pseudoword length (Fig 3A). Interactions were not significant (all *p*s > .478). Recognition accuracy did not differ between groups (main effect of group, *p* = .580), indicating comparable form-picture learning across TD and DLD. Repeated pseudowords were more accurately recognised than non-repeated ones (main effect of presentation type, β = 0.91, SE = 0.12, CI [0.66, 1.15], *p* < .001), and 2-syllable pseudowords were recognised more accurately than 4-syllable ones (main effect of pseudoword length, β = –0.61, SE = 0.12, CI [–0.85, –0.36], *p* < .001).

**Fig 3.**
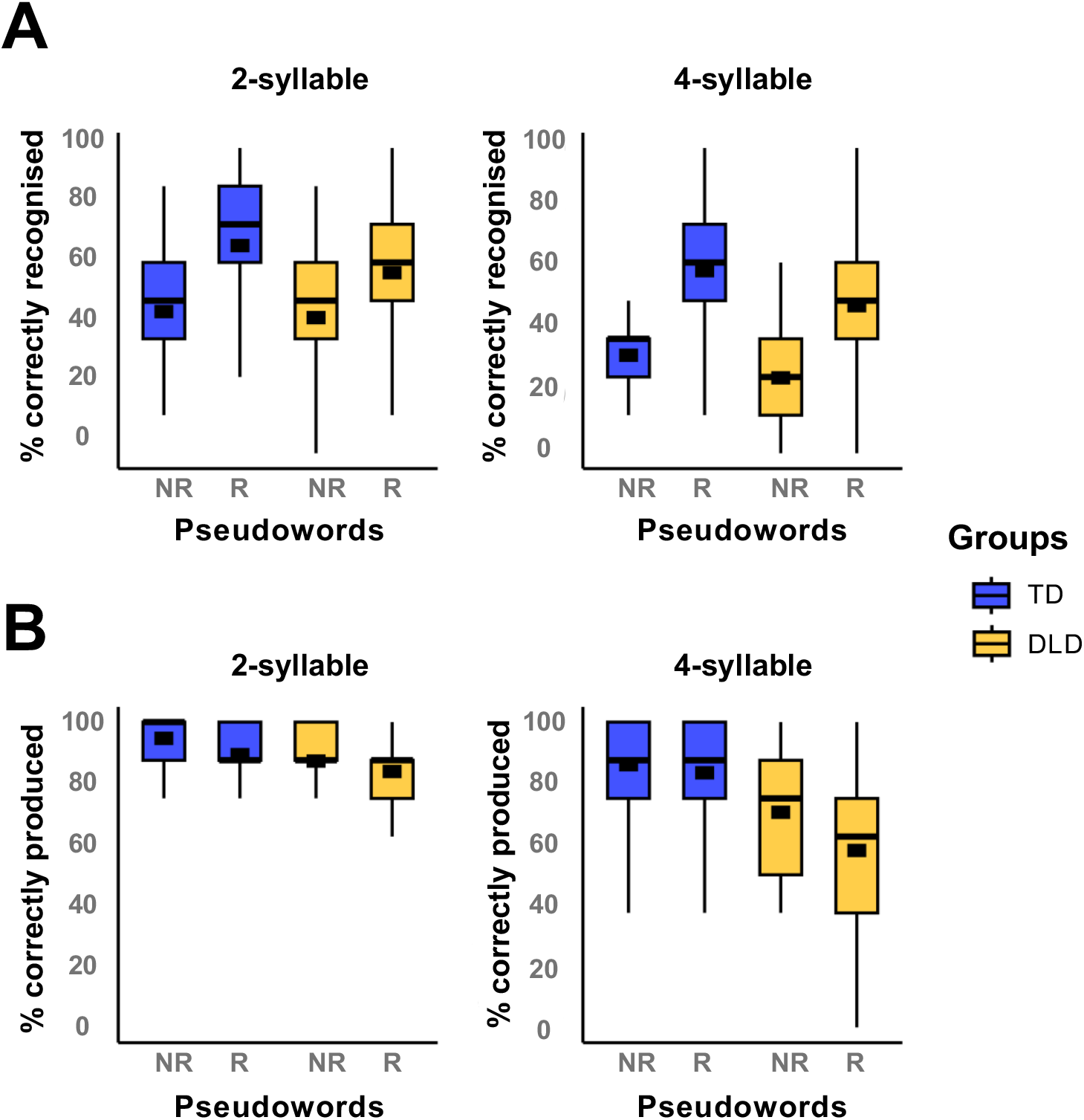
Behavioural performance during post-scan pseudoword recognition and repetition. (**A**) Percent correct form-referent recognition and (**B**) percent correct pseudoword repetition accuracy. Boxplots display the median and interquartile range (IQR), with whiskers representing 1.5 × IQR. Thicker black lines within each box denote group means. TD = typically developing; DLD = developmental language disorder; NR = non-repeated; R = repeated.

For post-scan repetition accuracy, a significant three-way interaction emerged between presentation type, group, and pseudoword length (β = –0.85, SE = 0.40, CI [–1.64, –0.07], *p* = .033; Fig 3B). While production accuracy was similar between groups on non-repeated 2-syllable pseudowords (*p*_adj_ = 1), the DLD group was less accurate than TD on repeated 2-syllable pseudowords (β = 0.93, SE = 0.27, CI [0.09, 1.77], *p*_adj_ = .022). For 4-syllable pseudowords, the DLD group showed lower production accuracy than TD on both non-repeated pseudowords (β = 0.95, SE = 0.21, CI [0.31, 1.58], *p*_adj_ = .001) and repeated pseudowords (β = 1.32, SE = 0.20, CI [0.72, 1.93], *p*_adj_ < .001).

### 3.2 Neuroimaging

#### Pseudoword repetition

In both TD and DLD groups, listening to and overtly repeating pseudowords evoked significant activity compared with baseline in a bilateral, symmetric network of regions encompassing the posterior inferior frontal cortex, lateral premotor cortex, sensorimotor cortex, supplementary motor area extending medially into the anterior cingulate gyrus, superior temporal gyrus, extending to mid-to posterior lateral parts, superior temporal sulcus, and large portions of occipital and inferior temporal cortex (Fig 4). Activity was more extensive in the left hemisphere in inferior frontal, inferior parietal and posterior temporal cortex, an asymmetry that was particularly evident in the TD group. Subcortically, activity was seen in the thalamus, amygdala, and globus pallidus bilaterally. Pseudoword repetition also evoked activity in cerebellar regions, including lobules V/VI anteriorly and lobule VIII and Crus I posteriorly.

**Fig 4.**
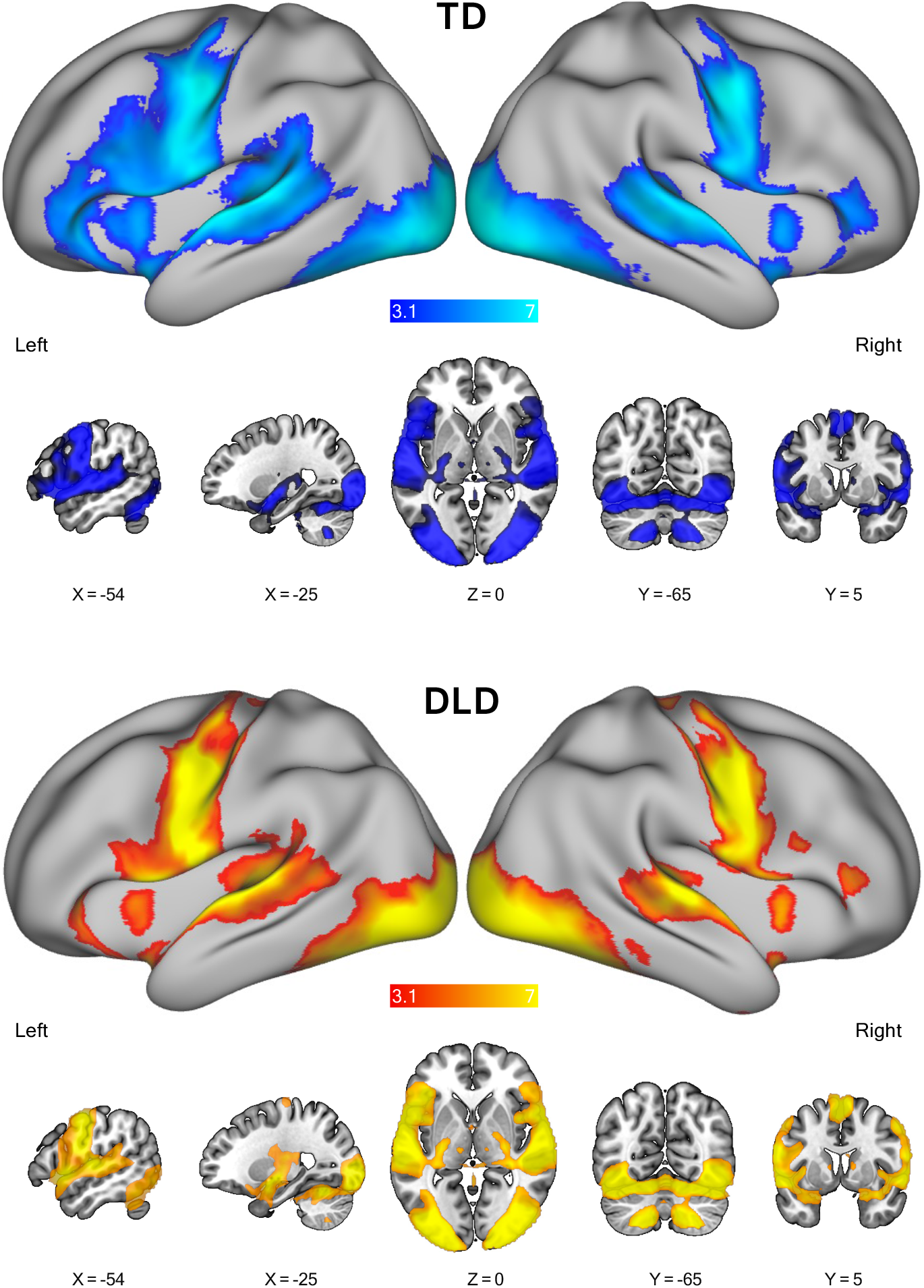
fMRI activity for pseudoword repetition per group. Statistical maps of mean BOLD activity are overlaid on the lateral views of inflated left and right hemispheres (top row) and the sagittal, axial, and coronal slices (bottom row). Coloured areas indicate significant clusters thresholded at Z > 3.1, *p* < .05 (corrected). Numbers below each slice indicate its position in mm relative to the horizontal and vertical planes through the anterior commissure.

The comparison between DLD and TD groups revealed significantly different levels of activity in lateral occipito-parietal cortex encompassing the posterior angular gyrus, medial parieto-occipital cortex encompassing retrosplenial cortex on the left, right posterior cingulate cortex, right dorsal post-central gyrus, left dorsal pre-central gyrus, and right superior frontal gyrus (Fig 5A and Table 1). Notably, none of these regions had activity that was significantly higher than baseline during the task, rather they showed task-negative activity that was lower than baseline or “deactivated”. The group differences were due to considerably greater task-negative BOLD responses in the TD group compared with the DLD group (Fig 5B).

**Fig 5.**
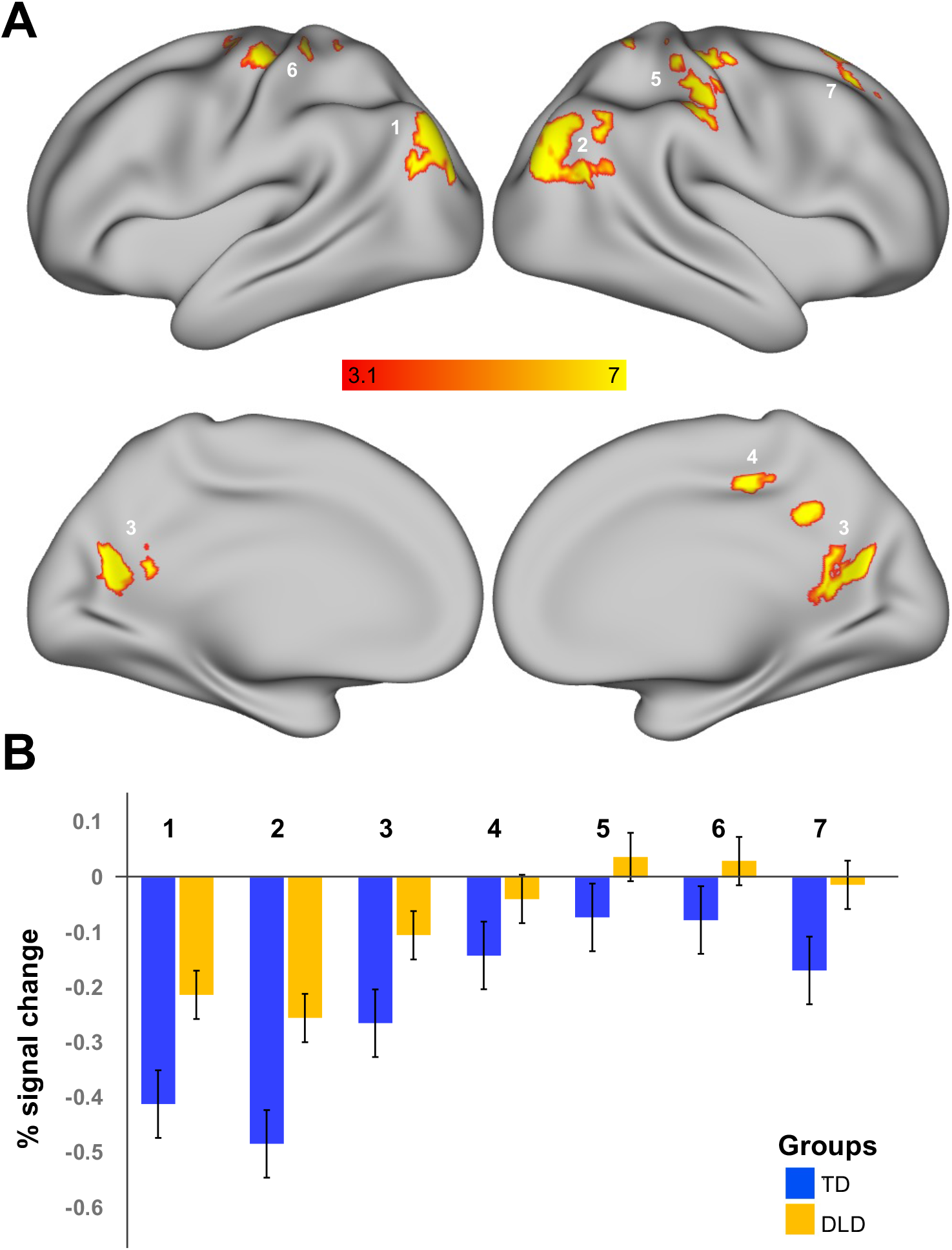
Brain regions where activity differed in TD and DLD groups during pseudoword repetition. (**A**) Statistical maps for the DLD > TD contrast overlaid on the lateral and medial views of inflated left and right hemispheres. Coloured areas indicate significant clusters thresholded at Z > 3.1, *p* < .05 (corrected). (**B**) The bar plots of percent signal change in each thresholded cluster from the DLD > TD contrast in seven regions presented for TD (blue) and DLD (yellow). Error bars represent standard error of mean (*SEM*). 1, 2 = L and R lateral occipito-parietal cortex; 3 = Medial parietal-occipital cortex; 4 = R posterior cingulate cortex; 5 = R dorsal postcentral gyrus; 6 = L dorsal precentral gyrus; 7 = R superior frontal cortex. See Table 1 and text for details.

**Table 1.** Group differences in brain activity during pseudoword repetition in DLD > TD contrast.

| Anatomical area | Coordinates |  |  | Z-statistic | Voxels |
| --- | --- | --- | --- | --- | --- |
|  | X | Y | Z |  |  |
| 1. L lateral occipital-parietal cortex | -42 | -82 | 22 | 3.91 | 443 |
| 2. R lateral occipital-parietal cortex | 44 | -78 | 30 | 4.71 | 997 |
| 3. Medial Parietal-occipital cortex | -16 | -62 | 18 | 4.09 | 346 |
| 4. R posterior cingulate cortex | 16 | -40 | 32 | 4.71 | 203 |
| 5. R dorsal postcentral gyrus | 30 | -32 | 58 | 4.05 | 556 |
| 6. L dorsal precentral gyrus | -34 | -16 | 58 | 4.33 | 431 |
| 7. R Superior frontal cortex | 18 | 18 | 58 | 4.00 | 224 |

*Note:* Contrast images for groups were thresholded at $Z > 3.1$ . Peak locations are presented for X (sagittal), Y (coronal) and Z (axial) coordinates in mm relative to the orthogonal planes through the anterior commissure, together with peak z-statistic and cluster extent in voxels. L - left; R - right.

The probabilistic overlap map for participants with DLD was largely like that for the TD group, but the proportion of each group showing activation of the same voxels was smaller in DLD in the superior temporal cortex bilaterally (Fig 6). This difference suggests less robust task-evoked activity in the DLD group due to greater spatial variability in the precise location activated, lower levels of activity at the statistical threshold used, or some combination of both effects.

**Fig 6.**
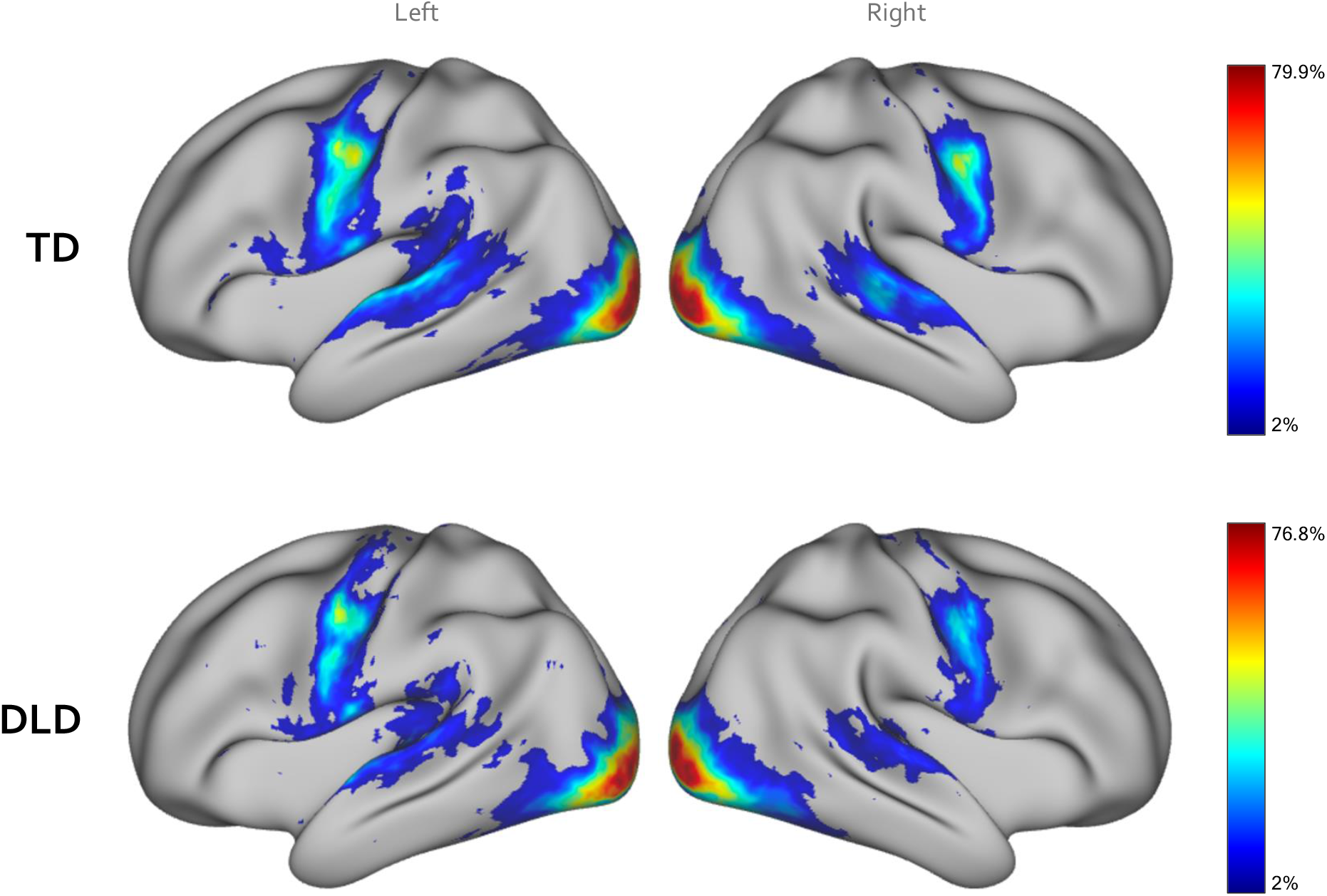
Probabilistic overlap maps for pseudoword repetition vs. rest contrast in TD and DLD groups. The maps depict the proportion of participants in each group (71 TD and 46 DLD) with significant activation in each voxel. Warmer colours illustrate areas of greater overlap.

#### Pseudoword learning

Brain activity in the TD group showed a linear decrease across the four presentations of pseudowords in regions encompassing the inferior frontal cortex, anterior insular, superior temporal and occipital cortex bilaterally (Fig 7). Medially, decreases in activity were visible in the supplementary motor area and anterior cingulate cortex. Activity in the prefrontal regions was strongly left lateralised. Decreases in activity were also seen in portions of the left and right cerebellar lobules (VIII). The statistical map showing learning-related linear decreases in activity in the DLD group was considerably less extensive than that seen in the TD group, encompassing portions of the left inferior frontal cortex, superior temporal cortex, and inferior occipital cortex bilaterally. Group differences did not survive statistical correction.

**Fig 7.**
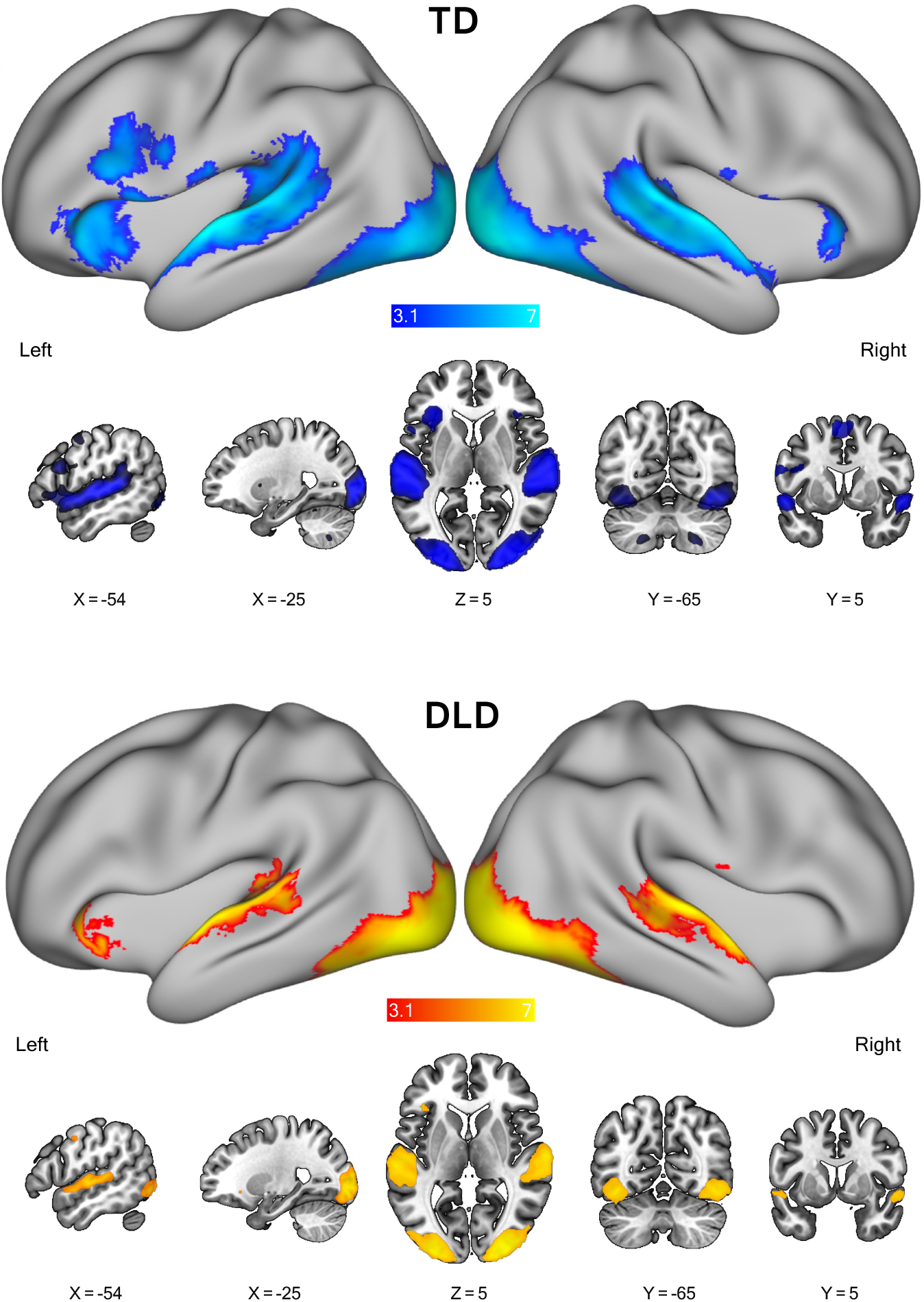
Brain regions showing learning-related activity decreases in TD and DLD groups. Statistical maps of areas showing this linear decrease are overlaid on the lateral views of inflated left and right hemispheres (top row) and the sagittal, axial, and coronal slices (bottom row). Coloured areas indicate significant clusters thresholded at Z > 3.1, *p* < .05 (corrected). Numbers below each slice indicate its position in mm relative to the horizontal and vertical planes through the anterior commissure.

#### ROI analysis

We extracted average percent signal change using anatomical masks from a frontal-striatal network involved in learning (pars triangularis, pars opercularis, supplementary motor area, caudate nucleus. and putamen), comparing activity across groups, hemispheres, and repetition trials. In the pars triangularis (Fig 8A), a significant group × hemisphere interaction indicated different lateralisation patterns in the TD and DLD groups (β = 0.09, SE = 0.03, CI [0.03, 0.11], *p* = .006); the TD group showed greater left-than right-hemisphere activity (β = 0.05, SE = 0.01, CI [0.03, 0.08], *p*_adj_ < .001), whereas the DLD group showed no hemispheric difference (*p*_adj_ = 1). Consistent with this, there was a main effect of hemisphere, with greater left-than right-hemisphere activity across groups (β = −0.05, SE = 0.02, CI [−0.11, −0.03], *p* = .007). No learning-related decreases were observed across repetitions (all *p*s > .08), and group differences averaged across hemispheres were marginal (*p* = .06).

**Fig 8.**
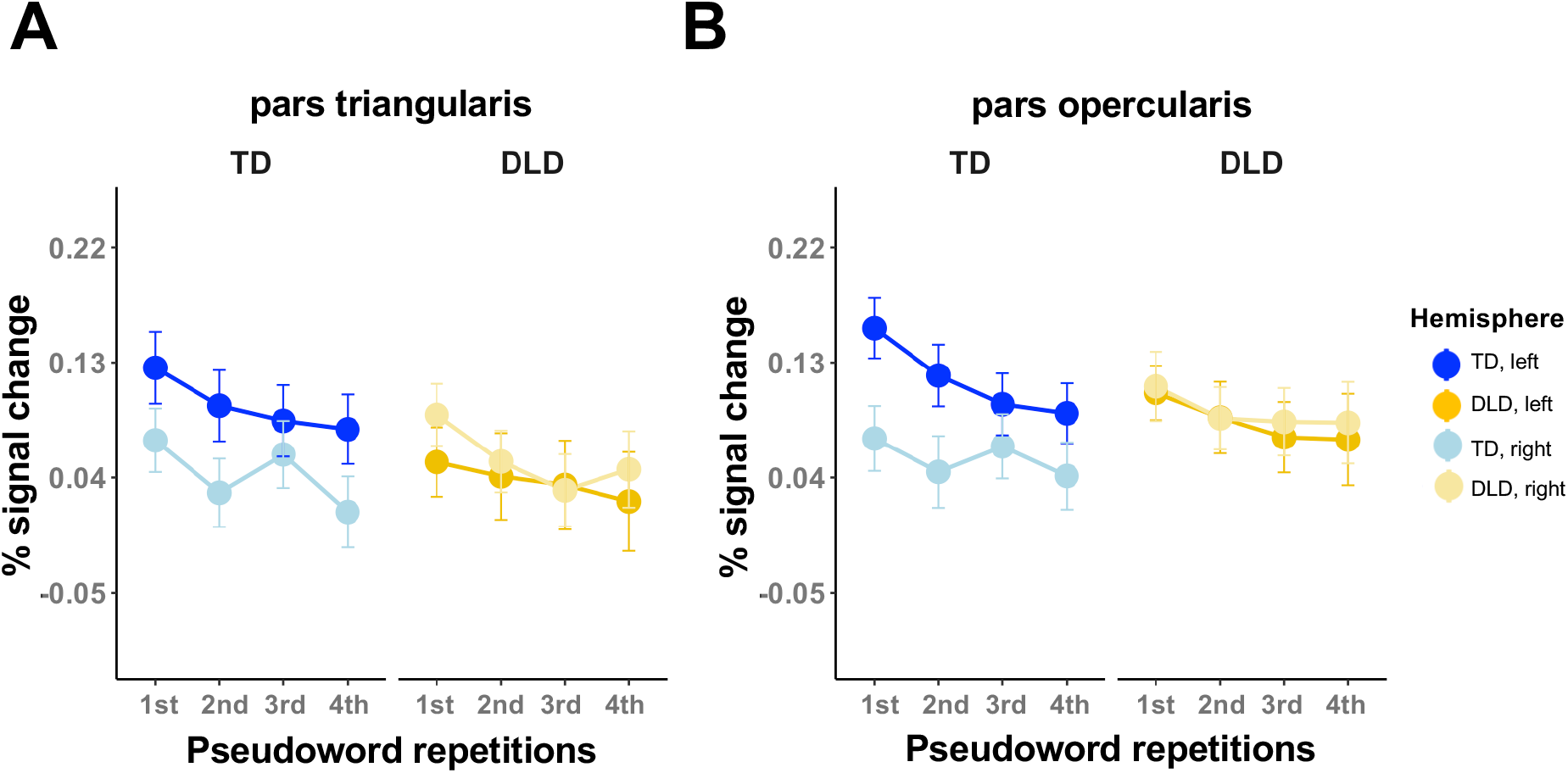
Brain activity associated with pseudoword learning in the inferior frontal gyrus in TD vs. DLD. The pattern of BOLD signal changes in pars triangularis (**A**) and pars opercularis (**B**) across the four pseudoword repetition trials in TD (blue) and DLD (yellow) groups. Darker and paler shades correspond to the left and right hemispheres, respectively. Group means are plotted. Error bars represent standard errors of the mean (*SEM*). left – left hemisphere; right – right hemisphere.

There was a similar group × hemisphere interaction in the pars opercularis (β = 0.09, SE = 0.02, CI [0.02, 0.19], *p* = .001; Fig 8B). The TD group again showed greater left-than right-hemisphere activity (β = 0.06, SE = 0.009, CI [0.03, 0.08], *p*_adj_ < .001), whereas the DLD group showed no hemispheric difference (*p*_adj_ = 1). A main effect of hemisphere indicated greater left-than right-hemisphere activity overall (β = −0.08, SE = 0.01, CI [−0.12, −0.05], *p* < .001). Across repetitions, significant learning-related decreases in activity were seen on the 3rd (β = −0.05, SE = 0.02, CI [−0.10, −0.01], *p* = .012) and 4th trials (β = −0.06, SE = 0.02, CI [−0.11, −0.01], *p* = .006). There was no main effect of group (*p* = .155).

There were no hemisphere, repetition, or group related differences in neural activity in the models for caudate nucleus, putamen, or supplementary motor area. Statistical results for these analyses are provided in Supplementary Tables 2-3.

## 4. DISCUSSION

Difficulties with pseudoword repetition are common in DLD, yet the neural mechanisms supporting pseudoword repetition and learning in this population are not well understood. Using fMRI, we examined whole-brain and frontostriatal activity during pseudoword repetition and learning in 46 participants with DLD and 71 typically developing (TD) controls aged 10– 15 years. Behaviourally, both groups showed learning effects during scanning, reflected through shorter duration of pseudoword productions over repetitions. Post-scan, form–referent learning was comparable across groups, whereas pseudoword repetition accuracy was lower in DLD, particularly for 4-syllable pseudowords. At the neural level, both groups engaged an extended network encompassing fronto-temporo-parietal regions (inferior frontal regions, premotor–SMA/anterior cingulate, sensorimotor cortex, and superior temporal regions), as well as occipital and inferior temporal areas. Activations in the TD were clearly left-lateralised in frontal and temporal areas as expected. Differences in activity were seen in portions of the default-mode network due to greater task-negative activity or deactivation in the TD group. Across repetitions, learning-related decreases in neural activity were observed in both groups in a left-lateralised inferior frontal areas and posterior temporal cortex bilaterally. Region-of-interest analyses confirmed the expected leftwards asymmetry in inferior frontal activation in the TD group but more bilateral recruitment in DLD.

### 4.1 Weaker task-related deactivation in DLD during pseudoword repetition

Significant group differences in the BOLD signal emerged in cortical regions where activation above baseline was not seen during pseudoword repetition. The differences in these regions instead reflected differences in task-induced deactivation. Specifically, participants with DLD showed significantly less deactivation below baseline compared with TD participants in the lateral occipito-parietal cortex encompassing the posterior angular gyrus, medial parieto-occipital cortex encompassing retrosplenial cortex on the left, right posterior cingulate cortex, right dorsal post-central gyrus, left dorsal pre-central gyrus, and right superior frontal gyrus. In TD participants, these regions were more robustly deactivated during task performance, whereas deactivation was markedly attenuated in DLD. These regions form core nodes of the default mode network (DMN), a network that is associated with coherent low-frequency activity at rest (Smith et al., 2009) and is reliably deactivated during cognitively demanding, goal-directed tasks (Raichle, 2015; Smallwood et al., 2021). We propose that reduced task-negative activity in the DLD group may reflect less efficient disengagement from the DMN during pseudoword repetition, which is arguably a linguistically demanding task for this population.

Additionally, because successful encoding depends on sustained attention to incoming stimuli (Erickson & Thiessen, 2015), weaker suppression of the DMN could plausibly interfere with attentional engagement during pseudoword repetition and learning (Buckner et al., 2013; Danckert & Merrifield, 2018). Converging evidence comes from resting-state work. Doucet et al. (2025) reported widespread connectivity differences in children with DLD across sensorimotor, cognitive-control, and default-mode networks. These findings fit with broader accounts that DLD involves cognitive differences that extend beyond language alone, including attention, executive functioning, and motor skills (Goffman & Gerken, 2023; Lukács et al., 2016; Sack et al., 2022; Tomas & Vissers, 2019). In particular, children with DLD often show weaker sustained attention in both auditory and visuospatial domains (Ebert & Kohnert, 2011), and auditory sustained attention has been linked to vocabulary knowledge in this population (Smolak et al., 2020), raising the possibility that reduced attentional engagement may contribute to difficulties in encoding and repeating unfamiliar phonological sequences. Caution is nonetheless required as these interpretations are tentative, especially given that our study was not intended to directly dissociate the language task from attentional demands.

### 4.2 Weak evidence for group differences in neural activity during pseudoword learning

In neurotypical adults, repetition-based pseudoword learning has been associated with linear reductions in BOLD signal across a largely left-lateralised auditory-motor network, thought to reflect increased neural efficiency with learning (Rauschecker et al., 2008). Broader theoretical accounts have additionally implicated corticostriatal circuits in the learning of phonological and speech-motor sequences (Krishnan et al., 2022; Ullman & Pierpont, 2005; Ullman et al., 2020), motivating our ROI analyses. We observed broadly similar whole-brain activation patterns in children. In TD participants, repeated pseudoword production was associated with reduced activity in regions implicated in speech articulation and phonological processing, including inferior frontal cortex, ventral premotor cortex, superior and middle temporal gyri, posterior superior temporal sulcus, and cerebellar lobules. Children with DLD likewise showed repetition-related decreases in these regions, although these effects were less extensive, particularly in left inferior frontal and anterior insular cortex. However, there were no robust whole-brain group differences in learning-related activity.

These findings do not support a broad difference in the neural systems underlying repetition-based pseudoword learning in DLD. Behaviourally, both groups showed faster pseudoword productions across repetitions, and neurally, both showed repetition-related decreases in activity at the whole-brain level. ROI analyses likewise provided no evidence of altered learning-related responses in the caudate, putamen, or supplementary motor area. They did, however, reveal reduced hemispheric specialisation of inferior frontal cortex in DLD: whereas TD children showed left-lateralised activation in the pars triangularis and pars opercularis during learning, children with DLD recruited left and right inferior frontal regions more similarly. This reduced lateralisation is consistent with previous observations in DLD and other neurodevelopmental disorders suggesting atypical hemispheric specialisation for language (Bishop, 1990; Xu et al., 2015). A potential explanation for this observation is that stronger left-lateralisation in the TD group reflects more efficient or specialised recruitment, whereas more bilateral engagement in DLD reflects either compensatory mechanisms (i.e., recruiting right homologues to assist in difficult tasks) or less specialised circuitry.

In sum, although the present results do not indicate a broad deficit in repetition-based implicit learning in DLD (to the extent that our fMRI design could be considered an implicit learning task), they remain consistent with some aspects of the Procedural (circuit) Deficit Hypothesis discussed in the introduction (Ullman & Pierpont, 2005; Ullman et al., 2020), insofar as the inferior frontal gyrus constitutes a key cortical node within frontostriatal circuitry. The reduced leftward specialisation of activity in DLD observed during learning, together with prior structural findings in this cohort from our group, points to neurobiological differences in this system in DLD. Specifically, Krishnan et al. (2022) reported altered myelin-related properties in dorsal striatal circuitry, whereas Bahar et al. (2024) found reduced surface area in inferior frontal cortex in the same cohort. Considered together with the present functional findings, these results suggest that DLD may involve converging structural and functional differences within a language-relevant frontostriatal network, potentially constraining the efficiency or specialisation with which it supports learning.

## Data and code accessibility

The anonymised neuropsychological and performance on pseudoword task and scripts supporting the findings presented are openly available on the Open Science Framework (https://osf.io/n76sz/overview?view_only=6f329bf021ab46409e9e62c34775b5a5).

## Author Contributions

**Nilgoun Bahar**: Conceptualization: Lead; Methodology: Lead; Formal analysis: Lead; Software: Lead; Investigation: Lead; Data curation: Lead; Visualization: Lead; Project administration: Lead; Writing – original draft: Lead; Writing – review & editing: Lead. **Gabriel J. Cler**: Methodology: Supporting; Formal analysis: Supporting; Software: Supporting; Investigation: Supporting; Data curation: Supporting; Resources: Supporting; Writing – review & editing: Supporting. **Salomi S. Asaridou**: Investigation: Supporting; Resources: Supporting; Project administration: Supporting; Writing – review & editing: Supporting. **Harriet J. Smith**: Methodology: Supporting; Investigation: Supporting; Data curation: Supporting; Project administration: Supporting; Writing – review & editing: Supporting. **Hanna E. Willis**: Methodology: Supporting; Formal analysis: Supporting; Investigation: Supporting; Data curation: Supporting; Validation: Supporting; Project administration: Supporting; Writing – review & editing: Supporting. **Sana Chughtai**: Data curation: Supporting. **Meron Haile**: Data curation: Supporting. **Máiréad P. Healy**: Investigation: Supporting; Resources: Supporting; Project administration: Supporting; Writing – review & editing: Supporting. **Saloni Krishnan**: Conceptualization: Lead; Methodology: Lead; Investigation: Lead; Data curation: Lead; Validation: Lead; Project administration: Lead; Formal analysis: Supporting; Funding acquisition: Supporting; Writing – review & editing: Supporting. **Kate E. Watkins**: Conceptualization: Lead; Methodology: Lead; Investigation: Lead; Data curation: Lead; Validation: Lead; Project administration: Supporting; Formal analysis: Supporting; Funding acquisition: Lead; Supervision: Lead; Writing – review & editing: Supporting.

## Acknowledgements

We are grateful to the children and families who generously participated in this research; their involvement made the study possible. We thank all the individuals and organisations who supported our recruitment efforts. We appreciate Professor Dorothy Bishop’s ongoing encouragement, and we thank Dr. Caroline Nettekoven for her contributions to data curation. We are indebted to colleagues at the Wellcome Centre for Integrative Neuroimaging, particularly the MRI team at the Oxford Centre for Human Brain Activity (Sebastian Rieger, Juliet Semple, Nicky Aikin, Nicola Filippini, Eniko Zsoldos, and Emily Hinson) for their technical expertise and support.

## Declaration of generative AI and AI-assisted technologies in the manuscript preparation process

During the preparation of this manuscript, the first author used OpenAI ChatGPT to refine the clarity and grammar of selected sentences. All content generated with AI assistance was reviewed, edited, and approved by the authors, who take full responsibility for the final manuscript.

