## Supplementary material for "Neural correlates of novel word-form learning in developmental language disorder": nwr_fmri_DLD_supplementary_biorxiv.pdf

[1] Department of Experimental Psychology & Oxford Centre for Integrative Neuroimaging, University of Oxford, Oxford, UK
[2] Department of Neurology, Dyslexia Center, University of California at San Francisco, San Francisco, USA
[3] Department of Speech & Hearing Sciences, University of Washington, Seattle, USA
[4] Division of Psychology and Language Sciences, University College London, London, UK
[5] Nuffield Department of Clinical Neuroscience, University of Oxford, Oxford, UK
[6] Department of Psychology, University of Cambridge, Cambridge, UK
[*] Joint senior authors

**Supplementary Table 1. Demographic and neuropsychological data for TD and DLD groups.**

| | TD (*n* = 71) | DLD (*n* = 46) | *p-value* | Group differences |
|---|---|---|---|---|
| **Demographics** | | | | |
| Mean age in years | 12.6 ± 0.2 | 12.4 ± 0.3 | 0.896 | --- |
| Age range (y;m) | 10;0–15;7 | 10;0–15;11 | --- | --- |
| Sex (M:F) | 40:31 | 32:14 | 0.214 | --- |
| Handedness (R:L) | 61:10 | 38:8 | 0.824 | --- |
| Language(s) spoken at home other than English (%Yes) | 13.84% (n = 65) | 4.34% | 0.116 | --- |
| **Language** | | | | |
| ROWPVT-4 | 128.7 ± 1.9 | 99.1 ± 2.3 | **< 0.001** | DLD < TD |
| EOWPVT-4 | 118.2 ± 1.8 | 91.1 ± 1.7 | **< 0.001** | DLD < TD |
| TROG-2 | 105.5 ± 1 | 81.3 ± 2 | **< 0.001** | DLD < TD |
| CELF-IV *Recalling Sentences*[1] | 11.8 ± 0.3 | 4.7 ± 0.4 | **< 0.001** | DLD < TD |
| ERRNI | | | | |
| *Initial Recall* | 100.4 ± 1.5 | 85.7 ± 1.8 | **< 0.001** | DLD < TD |
| *Delayed Recall* | 104.6 ± 1.3 | 86.6 ± 1.8 | **< 0.001** | DLD < TD |
| *Comprehension* | 107 ± 1.6 | 94.3 ± 2.2 | **< 0.001** | DLD < TD |
| NWR (max. 30)[2] | 26.3 ± 0.3 | 17.4 ± 0.8 | **< 0.001** | DLD < TD |
| **Memory** | | | | |
| CMS | | | | |
| *Digit Span Forwards*[1] | 11.6 ± 0.3 | 5.5 ± 0.4 | **< 0.001** | DLD < TD |
| *Digit Span Backwards*[1] | 11.8 ± 0.3 | 6.9 ± 0.5 | **< 0.001** | DLD < TD |
| *Word List Immediate Recall*[1] | 10.2 ± 0.4 | 6.1 ± 0.4 | **< 0.001** | DLD < TD |
| *Word List Delayed Recall*[1] | 10.3 ± 0.4 | 7.3 ± 0.5 | **< 0.001** | DLD < TD |
| *Word List Delayed Recognition*[1] | 8.6 ± 0.4 | 6.7 ± 0.5 | **0.004** | DLD < TD |
| **Reading** | | | | |
| TOWRE-2 | | | | |
| *Sight Word Efficiency* | 106 ± 1.3 | 81.8 ± 2 | **< 0.001** | DLD < TD |
| *Phonemic Decoding Efficiency* | 111.6 ± 1.6 | 80.8 ± 2.4 | **< 0.001** | DLD < TD |
| **Nonverbal reasoning** | | | | |
| WISC-IV | | | | |
| *Composite score* | 12.3 ± 0.2 | 8.6 ± 0.4 | **< 0.001** | DLD < TD |
| *Block Design*[1] | 13.3 ± 0.2 | 9.8 ± 0.4 | **< 0.001** | DLD < TD |
| *Matrix Reasoning*[1] | 11.3 ± 0.3 | 7.3 ± 0.5 | **< 0.001** | DLD < TD |
| *Coding*[1] | 9.6 ± 0.3 (n = 71) | 5.7 ± 0.4 | **< 0.001** | DLD < TD |
| **Motor** | | | | |
| NEPSY *Oromotor Sequencing* (max. 70)[2] | 60.5 ± 0.9 | 42.1 ± 1.6 | **< 0.001** | DLD < TD |
| Purdue Pegboard | | | | |
| *Dominant (z-score)* | -0.4 ± 0.1 | -1.5 ± 0.2 | **< 0.001** | DLD < TD |
| *Nondominant (z-score)* | -0.1 ± 0.1 | -1.1 ± 0.2 | **< 0.001** | DLD < TD |
| **Children's Communication Checklist-2** | | | | |
| GCC[3] | 84.7 ± 1.7 | 36.8 ± 3.1 (n = 45) | **< 0.001** | DLD < TD |

Note: Means ± SEM are provided unless otherwise stated. Sex, handedness ratios, and the results on the language background questionnaire were compared with a chi-squared test. All other variables were compared between groups using one-way analysis of variance (ANOVA), with post-hoc comparisons using Tukey's method. The last two columns display the p-values (p < 0.05 in bold typeface). Tests without a superscript denote standard scores with a standard mean of 100 and SD of 15.[1] Indicates scaled scores with a standard mean of 10 and SD of three. [2] Indicates raw scores. [3] Indicates scaled scores with a mean of 80 based on the sum of the eight subtests. M - male; F - female; R - right. L =-left. Vocabulary was assessed using the Receptive One-Word Picture

Vocabulary Test, Fourth Edition (ROWPVT-4; Martin & Brownell, 2011b) and the Expressive One-Word Picture Vocabulary Test, Fourth Edition (EOWPVT-4; Martin & Brownell, 2011a). Receptive grammar was assessed using the Test for Reception of Grammar, Second Edition (TROG-2; Bishop, 2003). Expressive grammar was indexed using the Recalling Sentences subtest of the Clinical Evaluation of Language Fundamentals, Fourth Edition (CELF-IV; Semel et al., 2003). Narrative abilities were assessed with the Expression, Reception and Recall of Narrative Instrument (ERRNI; Bishop, 2004).

**Supplementary Table 2. ROI analysis of neural activity during pseudoword learning in the caudate nucleus and putamen.**

| | Caudate nucleus | | | Putamen | | |
|---|---|---|---|---|---|---|
| **Predictor** | **β** | **SE** | ***p*-value** | **β** | **SE** | ***p*-value** |
| Repetition (1st) | | | | | | |
| 2nd | 0.032 | 0.02 | 0.117 | 0.008 | 0.02 | 0.664 |
| 3rd | -0.007 | 0.02 | 0.751 | -0.029 | 0.02 | 0.154 |
| 4th | -0.015 | 0.02 | 0.521 | -0.025 | 0.02 | 0.223 |
| Group (TD) | | | | | | |
| DLD | -0.003 | 0.03 | 0.915 | 0.035 | 0.03 | 0.339 |
| Hemisphere (left) | | | | | | |
| Right | 0.010 | 0.01 | 0.298 | 0.014 | 0.009 | 0.132 |
| Repetition trial × Group | | | | | | |
| 2nd × Group | -0.056 | 0.03 | 0.090 | -0.038 | 0.03 | 0.235 |
| 3rd × Group | -0.034 | 0.03 | 0.332 | -0.022 | 0.03 | 0.489 |
| 4th × Group | -0.004 | 0.03 | 0.897 | -0.009 | 0.03 | 0.764 |
| Repetition trial × Hemi | | | | | | |
| 2nd × Hemisphere | -0.004 | 0.01 | 0.740 | 0.001 | 0.01 | 0.9 |
| 3rd × Hemisphere | 0.011 | 0.01 | 0.431 | 0.014 | 0.01 | 0.306 |
| 4th × Hemisphere | -0.015 | 0.01 | 0.292 | -0.002 | 0.01 | 0.83 |
| Group × Hemisphere | 0.008 | 0.01 | 0.606 | -0.020 | 0.01 | 0.192 |
| 2nd × Group × Hemisphere | 0.001 | 0.02 | 0.965 | 0.009 | 0.02 | 0.665 |
| 3rd × Group × Hemisphere | -0.013 | 0.02 | 0.566 | 0.002 | 0.02 | 0.905 |
| 4th × Group × Hemisphere | 0.014 | 0.02 | 0.535 | 0.024 | 0.02 | 0.258 |

Note: Statistical results from linear mixed-effects models predicting neural activity for pseudoword learning based on repetition trial (1st, 2nd, 3rd, 4th), group (TD, DLD), hemisphere (left, right), and their interactions. Random intercepts were included for repetition trials and participants. The 1st trial, TD group, and left hemisphere were set as reference categories.

**Supplementary Table 3. ROI analysis of neural activity during pseudoword learning in the supplementary motor area**

| | Supplementary motor area | | |
|---|---|---|---|
| **Predictor** | **β** | **SE** | ***p*-value** |
| Repetition (1st) | | | |
| 2nd | -0.02 | 0.01 | 0.225 |
| 3rd | -0.03 | 0.01 | 0.043 |
| 4th | -0.02 | 0.01 | 0.248 |
| Group (TD) | | | |
| DLD | 0.02 | 0.03 | 0.514 |
| Repetition trial × Group | | | |
| 2nd × Group | 0.005 | 0.03 | 0.855 |
| 3rd × Group | 0.005 | 0.03 | 0.861 |
| 4th × Group | 0.008 | 0.03 | 0.787 |

Note: Statistical results from linear mixed-effects models predicting neural activity for pseudoword learning based on repetition trial (1st, 2nd, 3rd, 4th), group (TD, DLD), hemisphere (left, right), and their interactions. Random intercepts were included for repetition trials and participants. The 1st trial, TD group, and left hemisphere were set as reference categories.

### Appendix A

List of 32 pseudowords used in the fMRI study paradigm. The stimuli were created by the Wuggy Pseudoword Generator (Keuleers & Brysbaert, 2010). Half of the pseudowords were two syllables long and the other half contained four syllables. Each pseudoword was paired with the image of an alien during the in-scanner repetition task to assess children's learning of form-referent associations.

| **Non-repeated (NR)** | | **Repeated (R)** | |
|---|---|---|---|
| 2-syllable | 4-syllable | 2-syllable | 4-syllable |
| Samper | Belospofic | Leppy | Barmicory |
| Pargo | Mittedatic | Rantop | Fripacerus |
| Ruker | Bansenica | Banpet | Penicova |
| Demell | Matacolo | Meffon | Riscuplio |
| Fenter | Jaterbadon | Dombrin | Tosicotter |
| Henem | Impigatol | Tragle | Bodissary |
| Poyron | Vortenia | Zellop | Sycarery |
| Hombid | Fipotelar | Boosin | Cadesogist |
